# A human *ex vivo* model of radiation-induced skin injury reveals p53-driven DNA damage signaling and recapitulates a TGFβ fibrotic response

**DOI:** 10.1101/2025.06.04.657901

**Authors:** Caroline Dodson, Sophie M. Bilik, Gabrielle DiBartolomeo, Hannah Pachalis, Lindsey Siegfried, Jordan A.K. Johnson, Seth R. Thaller, Irena Pastar, Marjana Tomic-Canic, Anthony J. Griswold, Rivka C. Stone

**Author notes:** These authors contributed equally to this work (Caroline Dodson, Sophie M. Bilik).

## Abstract

Radiation-induced skin injury is a poorly understood complication affecting cancer patients who undergo radiotherapy, with no current therapies able to prevent or halt its progression to debilitating radiation-induced skin fibrosis (RISF). Addressing the need for clinically relevant human models, this study developed and characterized a human *ex vivo* skin model that recapitulates the temporal molecular processes of cutaneous radiation injury, as demonstrated through bulk RNA-sequencing and tissue validation studies. Human skin explants subjected to ionizing radiation demonstrated rapid induction of DNA double-strand breaks, followed by a robust, p53-driven transcriptional program involving genes related to cell cycle arrest, apoptosis, and senescence. Over time, the irradiated skin exhibited increasing activation of pro-fibrotic pathways, notably epithelial-mesenchymal transition and *TGFβ1*-mediated signaling. This resulted in upregulation of classic fibrosis markers such as *COL1A1*, *FN1*, and increased collagen thickness. Importantly, regulators of the p53 axis, MDM2 and miR*-*34a, was observed, implicating these factors as potential therapeutic targets to modulate the balance between repair of radiation injury and pathologic fibrosis. Transcriptome analysis of irradiated and non-irradiated breast skin from post-mastectomy patients showed notable concordance of p53 and pro-fibrotic gene signatures comparable to the *ex vivo* model, underscoring its translational relevance. This work provides a platform for identifying early biomarkers and testing therapeutic strategies to prevent or mitigate cutaneous radiation toxicities, including RISF, beginning with elucidating the dynamic interplay between the p53-mediated DNA damage response and the onset of fibrosis following radiation. Ultimately, this work aims to improve long-term skin health and quality of life for cancer patients.

**One Sentence Summary:** Human ex vivo skin recapitulates a clinically relevant p53-mediated DNA damage and pro-fibrotic response to radiation.

## INTRODUCTION

Radiotherapy is a critical component of cancer treatment that confers survival benefit for many common cancers, with at least 50% of oncology patients receiving ionizing radiation as part of their care (*1*). A significant consequence of radiotherapy is collateral damage to healthy tissues, particularly the skin, which lies in the path of the ionizing radiation beam as it travels to target and destroy tumor cells (*2, 3*). Approximately 95% of patients experience various skin reactions to radiotherapy, which range from mild to severe and are influenced by dose, overlapping fields, and treatment duration (*4*). While most acute reactions resolve within days to weeks, 30-70% of patients experience a continuous skin injury response which progresses to chronic debilitating radiation-induced skin fibrosis (RISF) resulting in disability, pain, and diminished quality of life in the months to years following radiation exposure (*4-6*). Current clinical management of cutaneous radiation toxicities focuses on symptomatic relief of acute skin reactions through supportive measures, such as topical emollients (e.g., aloe vera, Aquaphor), hyaluronic acid-based compounds, and corticosteroids in high-risk patients, as well as lifestyle changes like avoiding sun exposure and skin irritants (*7, 8*). However, no therapy has demonstrated efficacy in halting progression to RISF (*2*). Moreover, while animal models recapitulate some mechanistic and phenotypic features of RISF (*9*), translation of these findings to clinical therapies has not yet succeeded, underscoring the critical need for more translationally relevant systems. A deeper understanding of pathologic processes driving the evolution of RISF in *human* skin is needed to predict, mitigate, and treat skin fibrosis in patients receiving lifesaving radiotherapy.

As a self-renewing proliferative organ, the skin is particularly vulnerable to ionizing radiation, which disproportionately affects rapidly dividing cells (*10*). Specifically, basal keratinocytes, hair follicle stem cells, and melanocytes are highly radiosensitive, accounting for the alopecia and pigmentary skin changes that are commonly observed post-radiotherapy (*4, 11, 12*). Radiation-induced cellular damage initiates a cascade of events resembling phases of the classic wound healing response to barrier compromise (i.e., acute wounds), including inflammation, keratinocyte proliferation, and remodeling aimed at tissue repair (*12-14*).

One of the earliest and most critical events in the radiation-specific cascade is the DNA damage response, which is triggered by both direct and indirect mechanisms (*15*). Ionizing radiation directly causes DNA double-stranded breaks, leading to the activation of the ataxia telangiectasia mutated (*ATM*) kinase and subsequent phosphorylation of histone H2AX (γ-H2AX) (*16, 17*). Indirectly, radiation generates free radicals that contribute to additional DNA damage and oxidative stress (*10*). Among the key downstream effectors of *ATM* is the transcription factor *p53*, which regulates cell fate decisions following radiation injury — promoting cell cycle arrest to allow DNA repair, or initiating senescence or apoptosis when repair is unsuccessful (*17-21*). Fibrosis evolves post-radiation as it does in many other organs and tissues: unrepaired injury drives chronic inflammation that perpetuates *TGFβ* activation, collagen production, and excess extracellular matrix (ECM) deposition in the skin, leading to RISF (*2-4, 7, 22*). In this study, we present a human *ex vivo* model of radiation-induced injury designed to emulate the complex cutaneous response to ionizing radiation, spanning the continuum from acute injury and inflammation to progressive fibrosis. By highlighting its clinical implications through a comparative review with patient skin biopsies, this model acts as a reliable and translatable tool for investigating the primary, early drivers of RISF and for evaluating new therapeutic modalities. We focused on identifying and characterizing three elements underpinning this complex response: (1) the DNA damage response, (2) the role of p53 in modulating the radiation injury response, and (3) candidate pro-fibrotic drivers of skin fibrosis in other contexts, as they may serve as early biomarkers of evolving RISF.

## RESULTS

### Induction of DNA damage by ionizing radiation in human skin *ex vivo*

To investigate the molecular mechanisms driving the cutaneous radiation injury response, we developed a human *ex vivo* model in which skin obtained from patients undergoing elective panniculectomy procedures was irradiated with a single dose of 0 (non-irradiated), 3.5, or 6 gray (Gy) and maintained in culture at an air-liquid interface for 7 days. Non-irradiated skin from each donor was maintained under identical conditions and served as a control, minimizing inter-donor variability and ensuring observed changes are not attributable to being in culture alone (**Fig. 1A**). Irradiated and control skin from a total of 12 donors (**Fig 1B**) was collected immediately (30-60 min) and on days 1, 2, 4, 5 and 7 post-irradiation. We then profiled the cellular and molecular responses to radiation injury with a focus on epidermal keratinocytes, dermal fibroblasts and skin-resident immune cells.

**Fig. 1.**
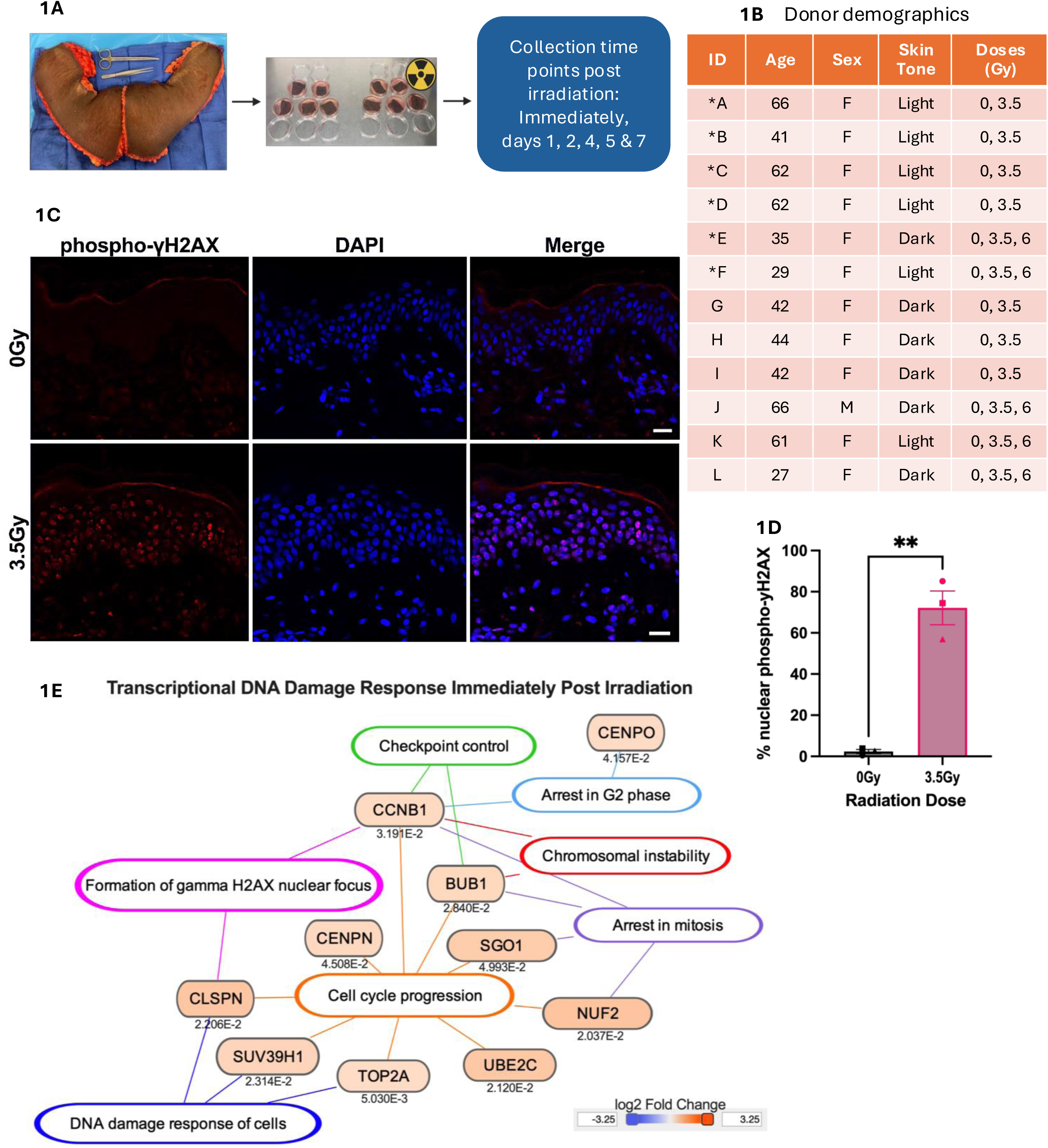
Establishment of a human ex vivo skin model and validation of early DNA damage responses to study radiation-induced injury. (A) Workflow schematic of the human *ex vivo* model used in this study (B) Donor demographics (ID, age and sex) and irradiation dose. Donors A-F, marked with an asterisk, were used for bulk RNA sequencing. (C) Representative immunofluorescence images at 1-hour post-irradiation showing phospho-γH2AX (red) as a marker of double-strand DNA breaks, DAPI (blue) nuclear counterstain, and merged images from irradiated (3.5 Gy) and control (non-irradiated) samples. (D) Quantification of phospho-γH2AX staining from (C), based on counts from three independent, blinded observers. Data represents N=3 donors; symbols correspond to individual donors; mean ± SD. Statistical significance was assessed using a paired t-test. (E) Ingenuity Pathway Analysis (IPA) of gene expression data in skin immediately post-irradiation, highlighting cell-survival-related genes and pathways. Differential expression was performed using edgeR on bulk RNA-seq data filtered for protein-coding genes. IPA Core Analysis was filtered to genes with p ≤ 0.05 and log_2_FC ≥ ± 0.5. Gene color reflects z-score; numerical values indicate Benjamini-Hochberg-corrected -log_10_ (p-value). Asterisks indicate statistical significance (* ρ < 0.05, ** ρ < 0.01).

As initial validation of our *ex vivo* model, we confirmed the induction of DNA double-strand breaks following irradiation via immunofluorescence (IF) staining for phospho-γ-H2AX, an established marker of DNA double-stranded breaks (*16, 17*). Irradiated skin displayed positive nuclear staining of γ-H2AX foci compared to controls immediately post-irradiation (**Fig. 1C-D**). This is largely resolved by 24 hours (1 day) post-irradiation, reflecting efficient repair of catastrophic DNA damage (*23*).

Having established the anticipated induction and repair of DNA damage following ionizing radiation, we proceeded to perform bulk RNA-sequencing on skin from six donors to characterize the broader, time-dependent radiation-induced response in our model. Transcriptomic profiles from irradiated (3.5 Gy) and control skin were compared at each timepoint: immediate (30-60 min), and days 1, 2, 5, 7 post-irradiation. Differentially expressed genes meeting significance thresholds of nominal p ≤ 0.05 and log2FC ≥ ± 0.5 (**Suppl Fig 1**) were subjected to Ingenuity Pathway Analysis (IPA), which applies Benjamini-Hochberg correction. Importantly, the early transcriptional signature immediately post-irradiation reflected the canonical acute DNA damage response (DDR; (*23*)), evidenced by significant upregulation of genes and pathways related to DNA damage, chromosomal instability, cell cycle arrest, and formation of γ-H2AX foci (**Fig. 1E**), consistent with our tissue staining in **Fig. 1C**. This initial validation substantiates that our *ex vivo* model replicates the initial radiation-induced molecular events as expected, thereby providing a robust platform for comprehensive mechanistic investigations into the progression of acute to chronic radiation injury.

### P53 drives the DNA damage response to ionizing radiation in human *ex vivo* skin

To further explore the radiation injury response in our *ex vivo* model over time, we utilized IPA’s Comparison Analysis tool, in which pathway analyses from days 1, 2, 5, and 7 (enrichment p-values and z-scores for pathways, processes and upstream regulators) were compared side-by-side and grouped by similarity (**Fig 2A**). The top common upstream regulator induced and activated across all days following irradiation was *TP53*. Other pathways and processes involving p53 signaling and cell death were similarly activated over time, while processes and regulators of cell cycle progression like Aurora kinases that control entry into mitosis (*24*) showed predicted inhibition. These findings support that our *ex vivo* model captures an early, robust, and sustained p53-mediated DDR to ionizing radiation. This is consistent with previous studies demonstrating the central role of p53 in driving the radiation response in both healthy and tumor tissues (*18-20, 25, 26*).

**Fig. 2.**
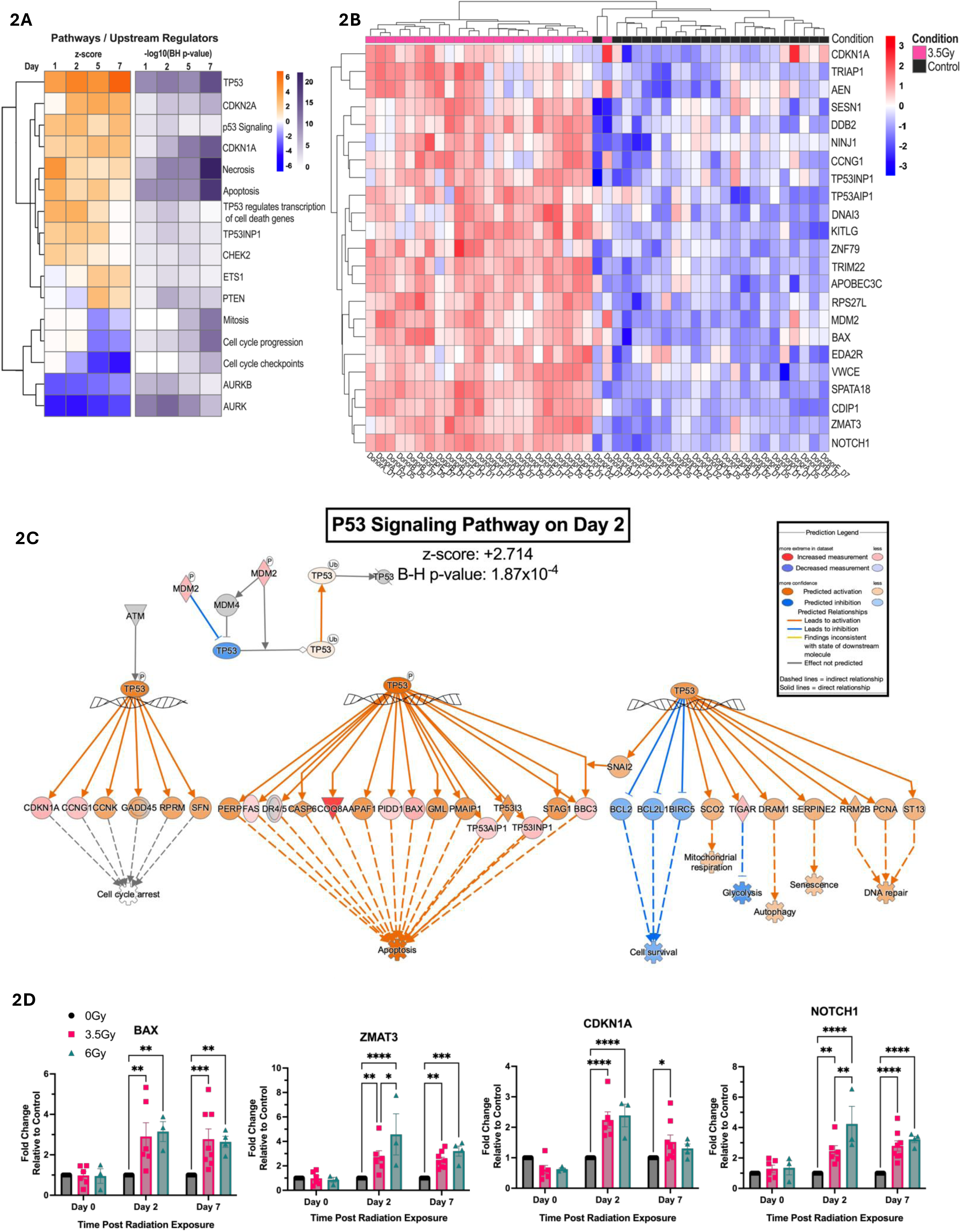
p53-Mediated DNA damage response in irradiated ex vivo skin. **(A)** Ingenuity Pathway Analysis (IPA) of predicted upstream regulators and biological pathways associated with DNA damage responses across days 1, 2, 5 and 7 post-irradiation. The first heatmap is colored by IPA z-score (predicted activation state), while the second shows Benjamini-Hochberg-corrected –log_10_ p-values (e.g., 1.3 corresponds to p ≤0.05). Differentially expressed genes were filtered for protein-coding, p ≤ 0.05, and log_2_FC ≥ ±0.5. Z-score heatmap rows were clustered by decreasing row average, with p-value heatmap rows ordered to match. **(B)** Heatmap of TP53-regulated genes from RNA-seq data, showing row-scaled log_2_-transformed CPM values. Red indicates increased expression; blue indicates decreased expression. Columns represent individual samples labeled by donor and time point (e.g., DonorA_D1, DonorB_D7). Annotation bars indicate condition above the heatmap (pink = 3.5Gy, black = control). **(C)** IPA-annotated “p53 signaling pathway” overlaid with RNA-seq gene expression changes at day 2 post-irradiation. Genes are colored by predicted activation state (orange = activated, blue = inhibited). **(D)** Quantitative PCR (qPCR) of selected p53 target genes in human *ex vivo* skin at days 0, 2 and 7 following 0 Gy (black, N=6), 3.5 Gy (pink, N=6) or 6 Gy (teal, N=3-4) radiation. Data were analyzed using the ΔΔCt method, normalized to *GAPDH*, and presented as fold change relative to matched 0 Gy controls for each donor and time point; mean ± SD. Statistical significance was assessed using a mixed-effects model with Tukey’s test for multiple comparisons. Asterisks indicate statistical significance (* ρ < 0.05, ** ρ < 0.01, *** ρ < 0.001, **** ρ <0.0001). All heatmaps were generated in RStudio using the pheatmap package.

We then visualized the expression patterns of *TP53*-regulated genes over time (**Fig. 2B**), revealing distinct clusters of upregulated genes whose induction was consistent across transcriptomic profiles of all six human donors. To visualize the functional context for these genes, we overlaid the post-irradiation day 2 profiles on IPA’s canonical “P53 signaling pathway” with associated enrichment p-value of 1.87x10^-4^ and activation z-score of +2.714 (**Fig. 2C**). In this annotated pathway, the directionality (up- or down-regulation) of the differentially expressed genes predicted activation of apoptosis, autophagy, senescence, and DNA repair processes, while cell survival and glycolysis/metabolism were predicted to be inhibited post-irradiation. Taken together, pathway analyses support that p53 is the core driver of the DDR to radiation in our *ex vivo* model; therefore, recapitulating findings from *in vivo* tissues (*19, 20, 27-30*). This further validates our human *ex vivo* model as a reliable platform for studying cutaneous radiation injury but also supports its utility for characterizing key cellular fates and testing interventions that modulate the p53 pathway.

To validate the RNA-seq findings, we selected a subset of p53 target genes with central roles in the canonical DDR: *BAX* and *ZMAT3* in apoptosis, *CDKN1A* (p21) in cell cycle arrest, and *NOTCH1* in p53 suppression (*19, 31, 32*). We explored the temporal and dose-dependent expressions of these genes in an expanded pool of eight donors treated with either 3.5 Gy or 6 Gy radiation. qPCR findings confirmed significant upregulation of these genes at day 2 and day 7 post-irradiation (**Fig. 2D, Suppl Fig 2**). We further observed that the 6 Gy dose elicited a more pronounced response compared to 3.5 Gy, and that *ZMAT3* and *NOTCH1* displayed a significant dose-dependent response two days post-irradiation. These findings further support the evolving temporal activation of p53 signaling and the induction of its downstream targets in our *ex vivo* human skin model.

### Autoregulation of the p53-mediated DDR by MDM2 and miR-34a-5p

We then shifted to explore regulators of the p53-mediated DDR in our model, which might serve as therapeutic targets to modulate the radiation injury response (*33*). *MDM2* is a well-established negative regulator of p53 signaling that undergoes post-translational modification followed by nuclear translocation during the p53-mediated DDR (*20, 34-36*). Bulk RNA-seq revealed differential expression of *MDM2* at 3.5 Gy, which was further supported by qPCR showing an early and dose-dependent induction of *MDM2* at day 2 across 3 radiation doses (0, 3.5 and 6 Gy; **Fig. 3A**). Western blot analysis identified an increase in MDM2 isoforms (55-90 kDa) at day 7 post-irradiation (**Fig. 3B & 3C**). DNA damage-induced phosphorylation of MDM2 at Ser395 by *ATM* kinase triggers its nuclear localization, where it binds and inhibits transactivation of p53 (*37*). Immunofluorescence staining using an anti-phospho serine 395-MDM2 antibody revealed MDM2 nuclear accumulation in irradiated tissue compared to non-irradiated controls (**Fig. 3D & E**).

**Fig. 3.**
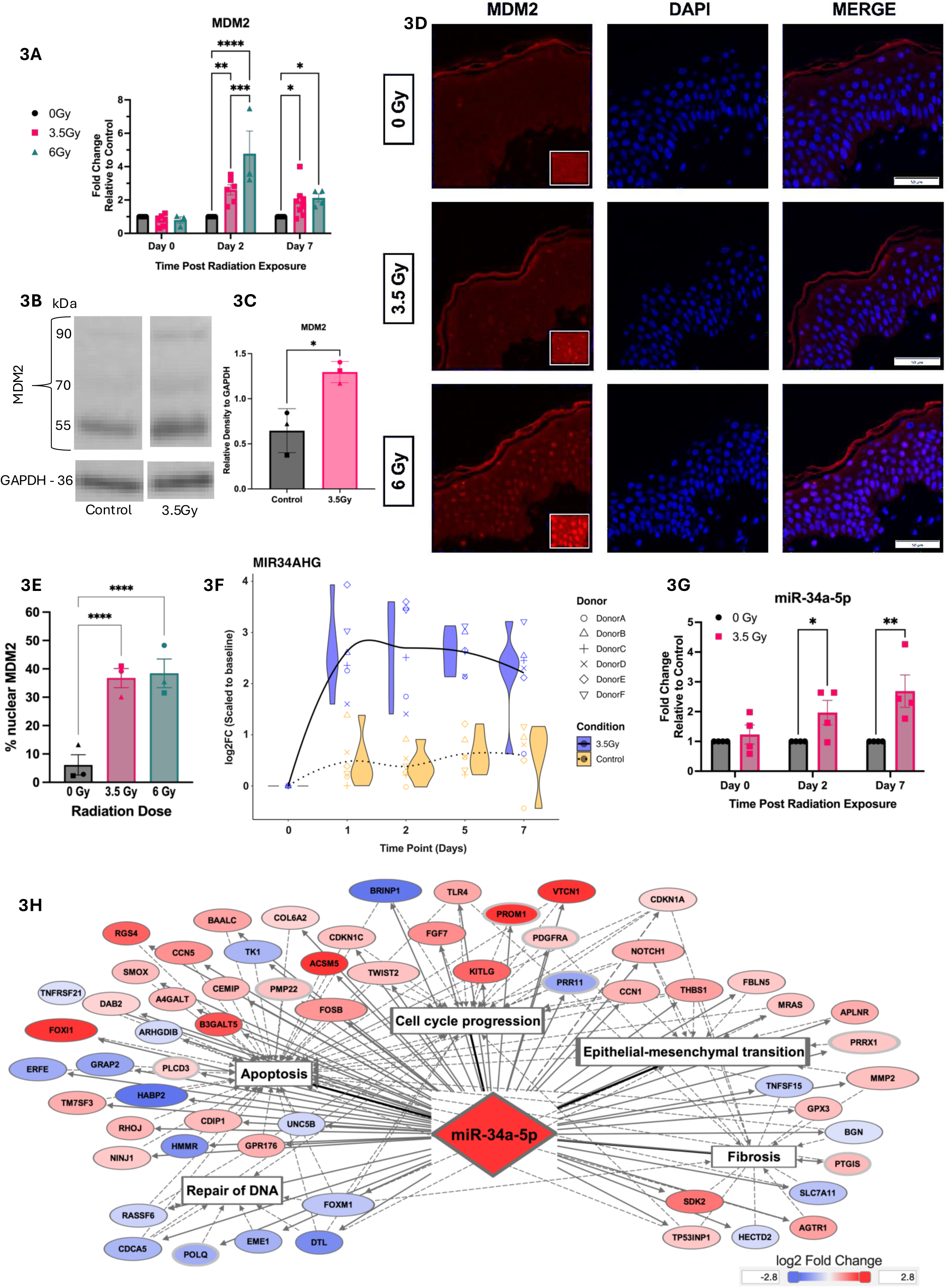
Regulators of p53-mediated DNA damage response. **(A)** Quantitative PCR (qPCR) of MDM2 expression in human *ex vivo* skin with 0 Gy (black, N=6), 3.5 Gy (pink, N=6) and 6 Gy (teal, N=3-4) at days 0, 2 and 7 post-irradiation. Expression was normalized to GAPDH, analyzed using the ΔΔCt method and presented as fold change relative to the 0 Gy control at each time point. **(B)** Representative western blot showing MDM2 protein expression at day 7 in control and 3.5 Gy samples. GAPDH served as the loading control. **(C)** Quantification of MDM2 protein levels **(B)**. Band intensities were extracted using ImageJ, and signals from multiple MDM2 isoforms (post-translationally modified) were averaged for each sample. Values were normalized to GAPDH. Each donor is represented by a unique symbol (N=3). **(D)** Representative immunofluorescence staining of phospho-MDM2 (Ser395) at day 1 post-irradiation, showing cytoplasmic localization in control samples and increased nuclear localization in irradiated samples (3.5 Gy and 6 Gy). **(E)** Quantification of phospho-MDM2 nuclear localization from **(D)**, expressed as the percentage of cells with nuclear localization. Data were collected by three independent, blinded observers and analyzed using two-way ANOVA with Tukey’s multiple comparisons test (N=3 donors; symbols correspond to individual donors. Donor shown in panel D is marked with a circle). **(F)** RNA-seq trajectory plot showing log_2_ fold change in MIR34AHG expression over time, scaled to the earliest time point. (G) qPCR analysis of mature *miR-34a-5p* expression using cDNA enriched for small RNAs. Expression was normalized to *SNORD48* and analyzed using the ΔΔCt method. Fold change was calculated relative to the 0 Gy control at each time point. **(H)** IPA generated network of genes regulated by *miR-34* at day 7 post-irradiation. Functional annotations highlight associations with apoptosis, cell cycle, fibrosis, EMT and DNA repair. Asterisks indicate statistical significance (* ρ<0.05, ** ρ<0.01, *** ρ<0.001, **** ρ<0.0001).

We also noted that *MIR34AHG,* the gene encoding microRNA (miR)-34a, was significantly upregulated in our RNA-seq dataset across donors and over time (**Fig. 3F**). *MiR34a* is a p53-induced non-coding RNA induced that sensitizes cells to p53-mediated apoptosis following radiation damage, thereby enhancing tissue radiosensitivity (*20, 38, 39*). It has also been identified as a potential target to protect tissue from radiation damage (*20, 39, 40*). Using a miR-specific qPCR assay, we validated the induction of *miR-34a-5p* post-irradiation, which was most significant at day 7 (**Fig. 3G**). To explore the functional consequences of *miR-34a* induction in our model, we used IPA microRNA Target Filter analysis to identify *miR-34a* targets among the genes regulated at day 7 post-irradiation in our RNA-seq dataset, and to determine the biological processes associated with these targets (**Fig 3H**). The resulting network encompassed DNA damage repair, apoptosis, and cell cycle regulation as anticipated, but also highlighted *miR-34a*’s role in the fibrotic response (*41*). Specifically, *miR-34a* target genes involved in epithelial-mesenchymal transition (EMT) and fibrosis were up- or down-regulated 7 days post-irradiation (**Fig 3H**), linking the early (p53-mediated DDR) and late (pro-fibrotic) radiation response in our *ex vivo* model. Collectively, these validation experiments underscore the exceptional accuracy of our *ex vivo* human skin model in recapitulating the dose- and time-dependent activation of p53 signaling and its key regulators and downstream targets following radiation. This robust and measurable response of human tissue positions our model as a valuable and predictive platform for the preclinical evaluation of cutaneous radiation injury mitigators and therapeutic strategies.

### Radiation triggers a pro-fibrotic response involving epithelial-mesenchymal transition

The second theme consistently maintained in the post-irradiation RNA-seq datasets after p53 signaling was fibrosis. Specifically, a series of pro-fibrotic upstream regulators and fibrogenic pathways were more significantly enriched and activated in day 7 vs. day 2 post-irradiation, suggesting a shift towards a more pro-fibrotic environment in irradiated skin over time. (**Fig 4A**). Among these, we noted predicted activation of EMT regulators SNAI1, SNAI2, and TWIST1; qPCR analysis confirmed significant transcriptional upregulation of *TWIST1* and *SNAI2* in our model 4 days post-irradiation (**Fig 4B**). Following skin injury, cutaneous EMT is marked by loss of E-cadherin and increased expression of vimentin as keratinocytes lose their cell-to-cell adhesion and acquire mesenchymal characteristics (*42*); therefore, we looked at these EMT mediators in irradiated *ex vivo* skin. We noted reduced expression of E-cadherin most prominently in the basal keratinocytes of irradiated skin, and total protein levels of E-cadherin decreased over time following irradiation (**Fig. 4C & 4D**). Conversely, the mesenchymal marker vimentin showed increased expression in irradiated dermis over time, indicating a shift towards a more mesenchymal phenotype (**Fig 4E & 4F**).

**Fig. 4.**
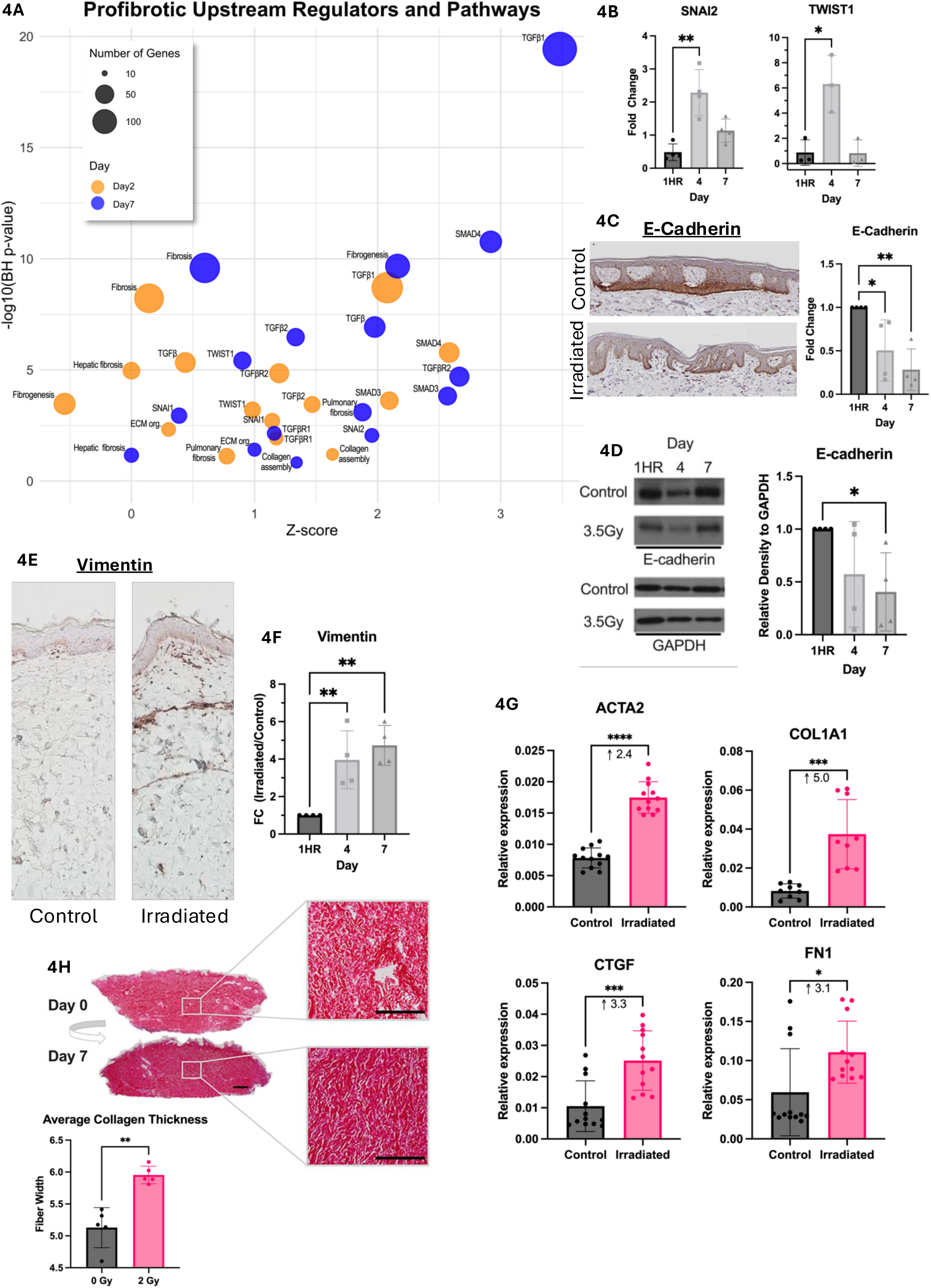
Induction of epithelial-mesenchymal transition (EMT) and pro-fibrotic genes in irradiated ex vivo skin. **(A)** Bubble plot showing enrichment of pro-fibrotic pathways, biological processes and upstream regulators at day 2 (yellow) and day 7 (blue) post-irradiation. The x-axis represents the predicted activation z-score, y-axis shows the Benjamini-Hochberg-corrected –log_10_ p-value, and bubble size reflects the number of overlapping genes. Data derived from IPA and plotted using ggplot2. **(B)** Quantitative PCR (qPCR) of EMT-inducing transcription in control (0 Gy) and 3.5 Gy-irradiated skin at hour 1, day 4 and day 7 post-irradiation. Expression was normalized to 18S and presented as fold change relative to control at each time point. **(C)** Representative IHC of E-cadherin in control and irradiated skin at day 7 post-irradiation. Quantification shown as fold change relative to control at each time point (mean ± SD; N=4). **(D)** Western blot of E-cadherin protein expression at 1 hour, day 4 and day 7 post-irradiation in control and 3.5 Gy samples. GAPDH was used as the loading control. Quantification of E-cadherin western blot shown as fold change relative to control at each time point (mean ± SD; N=3). **(E)** Representative immunohistochemistry (IHC) of vimentin in control and irradiated skin at day 7 post-irradiation. **(F)** Quantification of vimentin IHC over time, shown as fold change relative to control (N=4). **(G)** qPCR of pro-fibrotic genes in control and 3.5 Gy irradiated skin at day 7 post-irradiation. Expression was normalized to GAPDH and shown as relative expression (N=5). **(H)** Picrosirius red staining of 2 Gy-irradiated skin at day 0 and day 7 post-irradiation. Quantification of average collagen fiber thickness. Asterisks indicate statistical significance (* ρ < 0.05, ** ρ < 0.01, *** ρ < 0.001, **** ρ <0.0001). Statistical analyses: One-way ANOVA with Tukey’s correction for (B), (C), (D) and (F); two-tailed paired t-tests for (G) and (H).

### Induction of a *TGFβ1*-mediated fibrotic phenotype in *ex vivo* skin following radiation

Building on these early fibrotic signatures, our *ex vivo* model further demonstrated a robust *TGFβ1*-mediated fibrotic phenotype. *TGFβ1*, the master driver of tissue fibrosis (*43*), had the highest enrichment and activation scores among the pro-fibrotic factors with predicted involvement in the radiation response in our *ex vivo* skin, with over 100 of its downstream gene targets overlapping with the post-irradiation day 7 dataset (**Fig. 4A, Suppl Table 1**). To validate this in our model, we focused on the four genes previously identified as fibrotic *TGFβ1* response genes in human *ex vivo* skin and linked to excessive skin scarring: alpha smooth muscle actin *(ACTA2)*, type 1 collagen type (*COL1A1*), connective tissue growth factor (*CTGF*), and fibronectin 1 (*FN1*) (*44, 45*). We confirmed significant induction of all four genes post-irradiation (**Fig. 4G**), highest of which was a 5.0-fold increase in *COL1A1.* These effects persisted across radiation doses of 2-6 Gy (not shown). Furthermore, quantitative analysis of Picrosirius red-stained collagen fibers (*46*) identified a marked increase in average collagen thickness at day 7 in irradiated skin compared to controls (**Fig. 4H**). The successful induction and progression of these complex fibrotic changes, specifically EMT and collagen deposition, within our human *ex vivo* model highlights its substantial ability to serve as a clinically relevant platform for preclinical mechanistic studies of anti-fibrotic therapies for RISF.

### Biopsies of breast skin post-irradiation contain p53, inflammatory, and pro-fibrotic signals common to irradiated *ex vivo* skin

To directly assess the clinical-translational relevance of the radiation response identified in our *ex vivo* model, we analyzed RNA-seq profiles of *in vivo* irradiated and non-irradiated contralateral breast skin biopsies harvested from five post-mastectomy patients at the time of surgical reconstruction (GSE278183; **Fig. 5A**). Using IPA’s Comparison Analysis tool, we identified substantial overlap in pathways and biological processes between irradiated patient breast tissue and *ex vivo* irradiated skin. Among the top 25 shared upstream regulators, *TGFβ1* and *TP53* ranked highest based on the sum of BH-adjusted p-values across time points (**Fig. 5B**). Overlaying the gene datasets revealed 13 commonly regulated genes shared between the irradiated patient breast skin dataset and at least two time points of the irradiated *ex* vivo human skin dataset. These genes formed an interconnected network *involving TP53, TGFβ*, the matricellular protein SPARC (*47*), and *TNF* (**Fig. 5C**). Notably, commonly regulated *TP53* target genes in both irradiated patient breast biopsies and *ex vivo* skin included *MDM2* and *ZMAT3*, which we had validated in an expanded pool of *ex vivo* donors (**Figs. 2-3**). The p53-radiation response observed across all five patients was consistent across *ex vivo* donors and sustained over time (**Fig. 5D & 5E**). These compelling results offer substantial evidence that our human *ex vivo* skin model successfully characterizes the multifaceted and dynamic responses to radiation-induced injury — including the fundamental p53-mediated DDR and the progression towards fibrosis — making it a powerful and clinically relevant surrogate for investigating RISF and targeted therapeutic interventions.

**Fig. 5.**
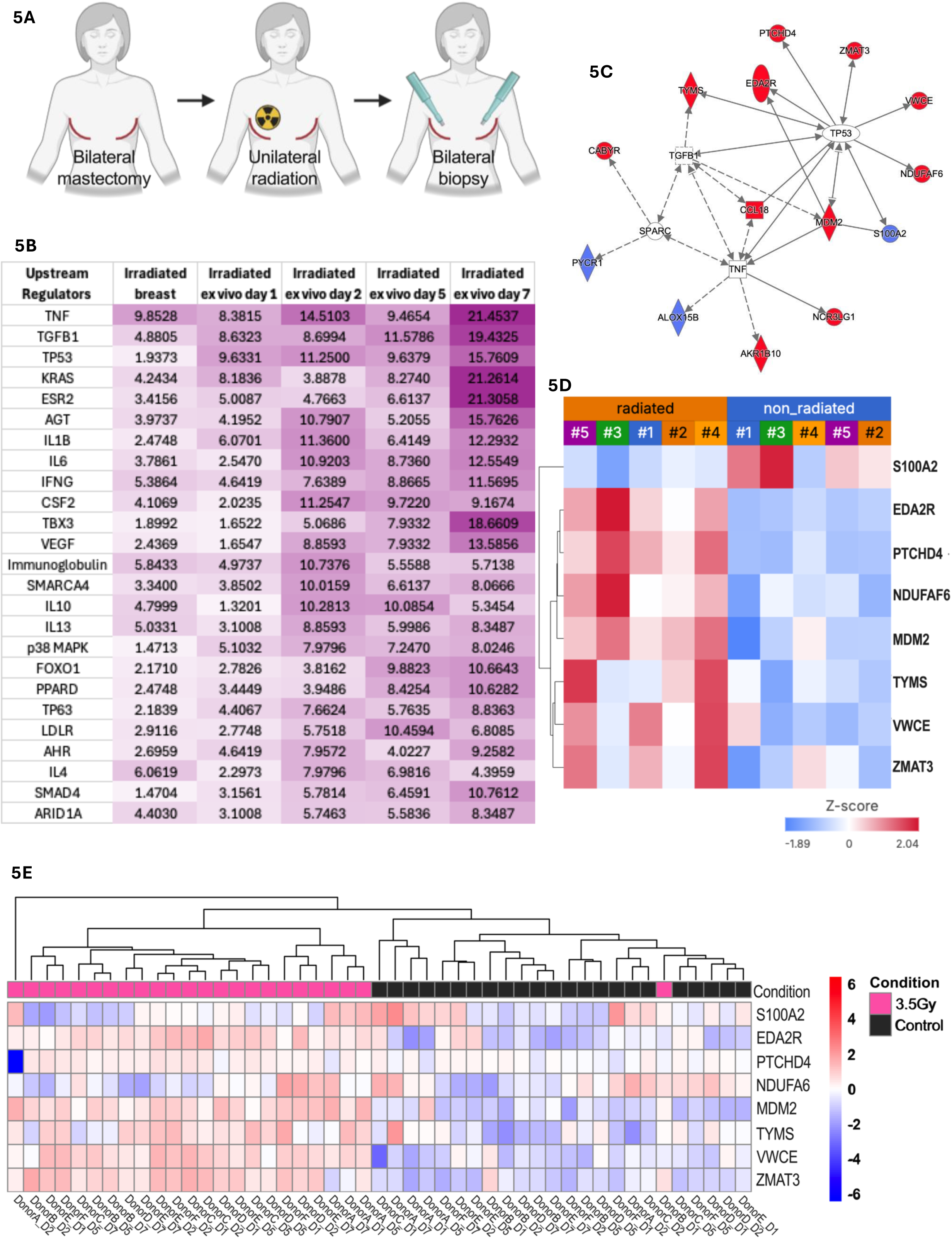
Correlation of ex vivo human skin radiation response with irradiated breast skin from patients post-radiotherapy. **(A)** Skin biopsies of the irradiated breast skin and non-irradiated skin of the contralateral breast were obtained from N=5 post-mastectomy patients at the time of surgical reconstruction. Raw bulk RNAseq profiles (GSE278183) were re-processed and re-analyzed using IPA. **(B)** Top 25 enriched upstream regulators common to irradiated breast skin and ex vivo irradiated skin across days 1, 2, 5, 7, ranked by sum of Benjamini-Hochberg (BH) enrichment p-values across all time points. Matrix values represent –log10(BH p-value). Colored by conditional formatting (0=white, max value=purple). **(C)** Network of the 15 genes overlapping between breast and at least 2 timepoints. Eight are downstream of *TP53* and 5 connect to *TGFβ1*/*SPARC* (role in ECM remodeling). Red indicates commonly upregulated; blue is commonly downregulated across datasets. **(D)** Heatmap of 8 commonly regulated *TP53* genes across profiles from non-irradiated and irradiated breast skin (patients numbered #1-#5). **(E)** Heatmap of the same *TP53* genes from panel **(D)** in human *ex vivo* skin. Data were log_2_-transformed counts per million (CPM) values. Red indicates increased expression; blue indicates decreased expression. Annotation bars indicate the condition above the heatmap (pink = 3.5 Gy, black = control). Heatmap was generated in Rstudio using the pheatmap package.

## DISCUSSION

While radiotherapy confers tremendous survival benefit for millions of cancer patients, it also produces a spectrum of cutaneous side effects, including severe skin fibrosis that significantly impairs quality of life of patients for months to years after radiation exposure. Despite this clinical burden, underlying mechanisms of radiation-induced skin injury and its evolution to fibrosis remain incompletely understood, hindering the development of effective therapies. To address this gap, we developed a human *ex vivo* model that demonstrates a reproducible temporal and dose-dependent radiation injury response, providing insights into the early evolution of radiation-induced skin injury and fibrosis. Importantly, our findings demonstrate that this model recapitulates key features of the *in vivo* radiation injury response observed in skin biopsies from post-radiotherapy surgical reconstruction patients, establishing its translational relevance.

We found that radiation rapidly initiates a DNA damage response centered on p53 and its downstream transcriptional program in our model, successfully recapitulating the acute radiation response observed in many organs and tissues *in vivo*. This pathway regulates cell fate decisions by inducing cell cycle arrest and senescence to allow DNA repair, or initiating apoptosis of irreparably damaged cells. The robust and expected activation of this fundamental cellular response in our *ex vivo* model further validates its ability to mimic the acute effects of radiation on human skin tissue. Along with p53 signaling, we identified a strong pro-fibrotic radiation response involving EMT and TGβ1 that mimics global phenotypic features of skin fibrosis. Fibrosis is a progressive process of abnormal matrix deposition that evolves following unresolved inflammation in the context of an ineffective healing response. Radiation injury exacerbates this process by further fueling the pro-inflammatory response and collagen production (*3*). While the canonical role of p53 in regulating the DNA damage/injury response to radiation is well-known (*19, 20, 25*), its causal role in regulating radiation-induced fibrosis in the skin and other organs has not been defined. Recent studies have noted p53’s pivotal role in biological processes that aggravate fibrosis, specifically TGFβ signaling, EMT, apoptosis, and senescence (*48-51*), supporting therapeutic targeting of p53 to manage hepatic, pulmonary, renal, and cardiac fibrosis (*52*). Importantly, not only did our *ex vivo* radiation injury model capture these pro-fibrotic biological processes (**Figs. 2-4**), but they were noted to be present *in vivo* in the irradiated breast skin from patients. This concordance between our model and clinical observations strongly suggests that sustained p53 signaling may be a critical driver in the transition from initial radiation injury to chronic fibrosis in the skin, highlighting the potential of our model to investigate this critical link.

We acknowledge the limitations of our study. *Ex vivo* skin is detached from the blood supply and does not capture the dynamic recruitment of immune cells and inflammatory mediators to the site of radiation injury. However, we have previously shown that resident immune cells, including Langerhans cells and gamma delta T-cells (*53-55*), are maintained in the *ex vivo* skin for at least 7 days using our culture conditions. Moreover, skin injury triggers the innate immune functions of keratinocytes, which release pro-inflammatory cytokines, produce anti-microbial peptides, and activate skin-resident immune cells. This may explain why the *ex vivo* skin successfully recapitulates many pro-inflammatory signals seen in the *in vivo* irradiated breast skin (**Fig. 5B**), despite its lack of blood supply. Additionally, cell populations within the skin exhibit differential radiosensitivity. However, bulk transcriptomic data make it difficult to resolve cell-specific signatures of radiation injury and fibrosis, potentially obscuring key radiation response pathways. Future studies leveraging spatial single-cell technologies are needed to further explore the cell-specific radiation responses, with the goal of targeting the cells driving a sustained pro-inflammatory and pro-fibrotic response to mitigate skin radiation injury.

Our findings suggest the connection between sustained p53-mediated DDR signaling and the pro-fibrotic skin response, though mechanistic studies are needed to validate a causal relationship. In addition to exploring this further in our model, linking sustained levels of p53 pathway biomarkers with clinical measures of evolving fibrosis in skin of patients in the days, months and years following radiotherapy will be a focus of future investigations. With regard to therapeutic interventions, our data conceptually support modulation of p53 regulators such as MDM2 and miR-34a to fine-tune the p53 response such that it acts to repair DNA, then shuts off before the progression of injury to debilitating fibrosis. For example, miR-34a suppresses EMT by targeting EMT-inducing transcription factors (*56*), so targeted modulation of its activity may mitigate the early pro-fibrotic response. Similarly, MDM2 inhibitors are currently in clinical development for cancer therapy and might be repurposed to manage cutaneous radiation toxicities.

In this regard, it is important to note that the p53-mediated effects are critically needed to induce death of cancerous cells within the tumor being targeted; as such, topical (rather than systemic) formulations are likely best-suited to mitigate and treat cutaneous radiation toxicities. However, even within the skin, transient p53 activity is temporarily needed to repair cells damaged by radiation as it passes through to reach the tumor. As such, therapeutic targeting would need to be timed to the inflection point at which p53 signaling exhausts the cell-reparative effects and shifts to drive excess pro-fibrotic signaling. Therefore, future studies aimed at identifying this critical time window and developing therapeutic strategies to precisely modulate p53’s activities following radiation injury are needed to prevent progression to severe skin fibrosis. Furthermore, better understanding of the early tissue biomarkers that best capture evolving RISF could serve as companion diagnostics to predict which patients are likely to progress to severe fibrosis, thereby enabling earlier intervention and alleviating suffering of patients experiencing impaired quality of life from cutaneous radiation toxicities.

## MATERIALS AND METHODS

### Study design

Human skin was obtained from subjects undergoing elective panniculectomy procedures under protocol approved by the University of Miami Institutional Review Board (IRB #20070922). Age, sex, and skin tone (e.g., light, medium or dark) of each subject was recorded. Specimens were otherwise de-identified and considered IRB-exempt as non-human subjects research under 45 CRF46.101.2 by the IRB at the University of Miami Miller School of Medicine. Specimens did not contain any of the 18 identifiers noted in the privacy rule, and therefore no informed consent was obtained.

Panniculectomy specimens that were < 6 in^2^, had visible striae involving >50% of the specimen, and/or had a severely thinned dermis, giving a more translucent appearance, were excluded from this study. Following excision in the operating room (OR), subcutaneous fat was removed from the skin samples, placed into sterile cups, and transferred into the biosafety cabinet. Each skin sample was examined in a 150 mm × 25 mm cell culture plate containing sterile phosphate-buffered saline (PBS) to assess tissue quality and ensure the removal of any remaining subcutaneous fat. Skin was then sectioned into approximately 2-inch by 2-inch pieces and subjected to a sequential decontamination process, via washes in five consecutive 50mL conical tubes containing 70% ethanol followed by three sterile PBS washes. After the final PBS wash, decontaminated skin sections were placed dermis-side down into fresh 150 mm × 25 mm TC plates containing pre-warmed, supplemented culture media. Media was prepared using 1X Dulbecco’s Modified Eagle Medium (DMEM) with GlutaMAX™ (Gibco, Cat. No. 10566016) as the base. To this, 10% non–heat-inactivated fetal bovine serum (FBS, Cytiva HyClone™, Cat. No. SH30071.03), 1% Antibiotic-Antimycotic (Gibco, Cat. No. 15240-062), and 1% sodium pyruvate (Gibco, Cat. No. 11360-070) were added. Media was thoroughly mixed by gentle inversion and vacuum filtered using a VWR® Bottle-Top Vacuum Filtration System with a 0.2 μm PES membrane (VWR, Cat. No. 514-0332) to ensure sterility. Each tissue culture plate received a sufficient amount of media to fully submerge the dermis without covering the epidermis and was placed on ice prior to further experimental procedures.

### Human ex-vivo RISF skin model and sample collection

Human skin specimens were irradiated with single doses of 3.5 or 6 Gy delivered by an Xstrahl cabinet irradiator, model RS225 (Xstrahl, Walsall, UK), with the following parameters: 190kV and 10mA with 0.5mm Cu filter for 3 minutes and 42 seconds for 3.5Gy or 6 minutes and 18 seconds for 6Gy. Irradiated skin was maintained with donor-matched (non-irradiated) control skin at an air-liquid interface, at 37°C/% CO_2_ for up to 7 days using a modified protocol from Mouawad et al. (*44*). Control and irradiated skin were collected at the following time points: immediate (30-60 min), and days 1, 2, 4, 5, and 7 post irradiation. At each time point, tissue samples were placed in RNAlater Stabilization Solution (Thermo Fischer Scientific, AM7020) for RNA isolation, snap-frozen for protein isolation, and fixed in 10% formalin for subsequent paraffin embedding.

### RNA Isolation and Quality Control

RNAlater-preserved samples were maintained on ice while being dissected and weighed (< 30mg). Samples were then finely chopped with a scalpel prior to being placed in tubes with RLT-βME with 3.0mm zirconium beads (Benchmark D10320-30). Samples then underwent 3-4 rounds of homogenization at 400 speed for 30 seconds with 1 minute rest between rounds in a BEADBug6 (Benchmark) homogenizer. Lysates were centrifuged at 16000 x g for 3 minutes, and the supernatant was thoroughly mixed with an equal volume of 70% ethanol. Total RNA was then isolated using QIAGEN’s RNeasy Mini Kit (QIAGEN Inc, Valencia, CA, USA) following manufacturer’s protocol. RNA concentration and purity was determined via NanoDrop spectrophotometry (NanoDrop 2000, Wilmington, DE) and RNA integrity was assessed via QIAxcel (Qiagen) to determine RNA Integrity Score (RIS). Samples with 260/230 > 1.6 and RIS> 4.5 were used for downstream studies.

### Quantitative Real-Time qPCR

1.0ug of total RNA from *ex-vivo* sample were reverse transcribed using QuantiTect Reverse Transcription Kit (Qiagen, Hilden, Germany) to generated cDNA, and real-time qPCR was performed in triplicate using PerfeCTa SYBR Green Supermix (QuantaBio, Beverly, MA) and the Bio-Rad CFX Connect system. The relative expression of target genes was normalized to housekeeping genes *18S*, *ARPC2* or *GAPDH* (or a combination thereof) based on the requirements of each qPCR analysis. For microRNA (*miR-34a-5p*) qPCR, 50-75ng of total RNA from *ex vivo* samples was reverse transcribed and enriched for microRNAs using microScript microRNA cDNA Synthesis Kit (Norgen Biotek, Ontario, Canada). Real-time qPCR was performed as stated above with relative expression normalized to levels of SNORD48. Primer sequences can be found in **Supplementary Table 2**.

### RNA sequencing

RNA from six donors underwent ribosomal RNA depletion with the resulting mRNA used for library preparation and subsequent bulk RNA-sequencing on the Illumina NovaSeq X Plus at Azenta Life Sciences (New Jersey, USA). Library preparation was performed by Azenta Life Sciences using standard RNA-sequencing protocol, yielding an average of 25-30 million paired-end (PE) reads per sample. Quality control of the raw sequencing reads was initially assessed using FastQC (*57*). Adapter sequences and low-quality bases were trimmed using TrimGalore in paired-end mode (*58, 59*). Trimmed reads were then aligned to the GRCh38 human genome using the STAR aligner (version 2.7.10a) with gene counts being quantified from GENCODE Release 44 gene annotations (*60*). Raw data files and the associated count matrix are available at NCBI GEO accession no. GSE297035. Alignment metrics were generated using GATK (version 4.3.0.0) CollectRnaSeqMetrics tool, with the hg38 reference file for annotation (*61*). MultiQC visually summarized all previous steps (*62*). Batch correction was performed using ComBat (version, 3.54.0; (*63*)) prior to paired differential expression via edgeR (*63, 64*). Analyses were restricted to protein-coding genes, and genes were considered differentially expressed if they met a nominal p ≤ 0.05 and log_2_FC ≥ ± 0.5. This threshold was chosen to balance specificity and sensitivity in identifying time-dependent gene expression changes across donors, and to enable robust downstream exploratory pathway analysis using Ingenuity Pathway Analysis (IPA), which applies Benjamini-Hochberg correction to account for multiple testing.

### Bioinformatics and Pathway Analyses

All RNA-seq data processing and visualization were performed in R (2024.12.1+563)(*65*). Visualizations were generated using packages including edgeR, tidyverse (e.g., dplyr, ggplot2, tidyr) (*66*), RcolorBrewer (*67*), pheatmap (*68*), gplots (*69*), and gridExtra (*70*). Heatmaps were scaled by row and clustered using Euclidean distance unless otherwise noted. Bubble plot was created with ggplot2.

Differentially expressed genes (DEGs) identified at each time point (3.5 Gy vs. control) underwent Ingenuity Pathway Analysis (IPA, Qiagen) Core Analysis which yielded enrichment p-values and predicted activation (z-scores) for canonical pathways, biological processes and upstream regulators. Enrichment p-values were corrected for multiple testing using the Benjamini–Hochberg method, and results with BH-adjusted p ≤ 0.05 and an absolute z-score ≥ 2 were considered significant.

### Total Protein Isolation and Western blotting

For Western blotting, 15–25 mg of snap-frozen tissue was lysed in RIPA buffer containing 150 mM NaCl, 1% NP-40/Triton X-100, 0.5% sodium deoxycholate, 0.1% SDS, and 50 mM Tris (pH 8.0), supplemented with Protease and Phosphatase Inhibitor Cocktail (Benchmark Scientific, Inc., Sayreville, NJ, USA). Tissue homogenization was performed using 3 mm triple-pure zirconium beads (Benchmark Scientific, Inc.) in a BeadBug microtube homogenizer (Benchmark Scientific, Inc.). Protein concentration was quantified using the Qubit™ Protein Assay Kit (Thermo Fisher Scientific, Cat. No. Q33211) and the Qubit 4 Fluorometer (Thermo Fisher Scientific, Walton, MA, USA).

Equal amounts of total protein were resolved by SDS-PAGE using 4–20% Criterion™ TGX™ Precast Gels (Bio-Rad), followed by transfer to polyvinylidene difluoride (PVDF) membranes (Bio-Rad). Membranes were blocked in TBST (TBS + 0.1% Tween-20) containing 5% non-fat dry milk (NFDM) for 1 hour at room temperature and then incubated overnight at 4°C with primary antibodies diluted in blocking buffer. Membranes were washed in TBST and incubated with HRP-conjugated secondary antibodies for 1 hour at room temperature. Bands were visualized using chemiluminescent substrate and imaged using a Bio-Rad ChemiDoc™ MP Imaging System. Band intensities were quantified using ImageJ (NIH) by background-subtracted densitometry and normalized to corresponding loading controls. Antibodies, including vendor, catalogue number and dilution are listed in **Supplementary Table 3**.

### Tissue immunostaining

Formalin-fixed paraffin-embedded (FFPE) tissues were sectioned at 5-6 μm using a microtome, deparaffinized in xylene (EMD Millipore, Gibbstown, NJ, USA), and rehydrated through a graded ethanol series. Antigen retrieval was performed by incubating slides in sodium citrate buffer (10 mM sodium citrate, 0.05% Tween-20, pH 6.0) at 95°C for 30-45 minutes, followed by a 20-minute cool-down at room temperature for IHC and an ice bath for IF. Endogenous peroxidase activity was quenched (for IHC only) using 0.3% hydrogen peroxide in methanol. Slides were washed with TBST (Tris-buffered saline, 0.1% Tween-20) and blocked for 1 hour at room temperature with 2% normal goat serum (Sigma-Aldrich, St. Louis, MO, USA) in TBST or 10 minutes with Background Punisher (BIOCARE Medical, Pacheco, CA, USA).

Primary antibodies were diluted in SignalStain® Antibody Diluent (Cell Signaling Technology, Danvers, MA, USA) for IHC and 5% BSA/TBST for IF, then applied overnight at 4°C. For immunohistochemistry (IHC), detection was performed using SignalStain® Boost IHC Detection Reagent (HRP, Rabbit) and developed using SignalStain® DAB Chromogen (Cell Signaling Technology). Slides were counterstained with Harris hematoxylin (Leica Microsystems, Wetzlar, Germany), dehydrated through ethanol and xylene, and mounted using Cytoseal™ XYL (Epredia, Cat. No. 8310-4). For immunofluorescence (IF), after incubation with primary antibodies and washing, sections were incubated with fluorophore-conjugated secondary antibodies (Alexa Fluor 488 or 594; Invitrogen, Carlsbad, CA, USA) for 1 hour at room temperature. Nuclei were counterstained with DAPI and slides were mounted with antifade mounting medium. IHC slides were imaged using the Olympus VS200 Slide Scanner and analyzed in QuPath v0.5.1 for percent positive area, staining intensity, and area fraction. IF slides were imaged using the Olympus DP74 Confocal.

All antibodies used, including vendor, catalog number, and dilution, are listed in **Supplementary Table 3**.

### Quantification of tissue immunostains

Quantification of IHC staining (E-cadherin and vimentin) was performed using QuPath v0.5.1 as previously described (*71*). Positive staining was normalized to the percent positive area per 100,000 μm² tissue section. Quantification of IF staining (γH2AX and MDM2) used the percentage of positive nuclear staining as manually scored by three independent, blinded observers, calculated as 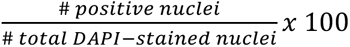, and the average values were used for analysis.

### Assessment of Dermal Collagen Thickness

Picrosirius Red staining was performed using the Picrosirius Red Stain Kit (Polysciences, Inc., Cat. No. 24901) to visualize and quantify collagen types I and III in paraffin-embedded tissue sections. Sections were deparaffinized in xylene and rehydrated through a graded ethanol series finishing with distilled water. Nuclei were stained with Weigert’s hematoxylin for 8 minutes, followed by thorough rinsing in distilled water. Slides were then stained in Solution A for 2 minutes, rinsed in distilled water, and subsequently stained in Solution B for 60 minutes. After staining, sections were placed in Solution C for 2 minutes, followed by a brief rinse in 70% ethanol for 45 seconds. Slides were then dehydrated through graded ethanol, cleared in xylene, and mounted using a suitable mounting medium. Stained sections were imaged using a polarized light microscope with multiple fields of view being imaged to cover representative areas of each tissue section. Collagen fiber analysis was conducted using the CT-FIRE software, as previously described (*46*), where images of Picrosirius Red-stained sections were uploaded to the CT-FIRE software for segmentation and quantification of collagen fibers, providing metrics on collagen fiber thickness.

### Statistical analyses

Data were analyzed using Prism software (GraphPad, La Jolla, CA, USA) and R (version 2024.12.1+563). For RNA-seq analyses, differential gene expression was assessed using edgeR with a paired design to account for matched control and irradiated samples from the same donor. P-values were adjusted for multiple testing using the Benjamini-Hochberg method, and genes with an adjusted p-value ≤ 0.05 and an absolute log_2_ fold change ≥ ± 0.5 were considered differentially expressed.

For qPCR experiments, relative gene expression was calculated using the ΔΔCt method. At each time point, irradiated samples were normalized to corresponding controls to determine fold change. Statistical testing was performed using a mixed-effects model with a repeated measures design using Tukey’s test for multiple corrections for within-time-point multiple comparisons (GraphPad Prism).

## ACKNOWLEDGEMENTS

The authors gratefully acknowledge Dr. Carol Feghali-Bostwick of the Medical University of South Carolina for her insights regarding the development of the *ex vivo* model, and Dr. Stuart Samuels of the Department of Radiation Oncology at University of Miami for his insights regarding translational applicability of radiation dosimetry and findings of the *ex vivo* model.

## FUNDING

Dermatology Foundation Physician Scientist Career Development Award (to RCS) National Institutes of Health grant R61/R33 DK131897 National Institutes of Health grant R01AR083385 (IP and MTC)

## AUTHOR CONTRIBUTIONS

Conceptualization: RCS, MTC, IP, AJG

Methodology: GD, LS, RCS, SRT, AJG, CD, SMB

Investigation: CD, SMB, GD, HP, LS, JAKJ

Visualization: CD, SMB, GD, HP, RCS, AJG

Funding acquisition: RCS, MTC, IP

Supervision: RCS, AJG, MTC, IP, SRT

Writing – original draft: CD, SMB, RCS

Writing – review & editing: CD, SMB, GD, HP, LS, JAKJ, SRT, IP, MTC, AJG, RCS

## COMPETING INTERESTS

None

## DATA MATERIALS AND AVAILABILTY

RNA sequencing data (raw files and count matrix) have been deposited to NCBI Gene Expression Omnibus under accession no. GSE297035.

## LIST OF SUPPLEMENTARY MATERIALS

**Fig. S1.**
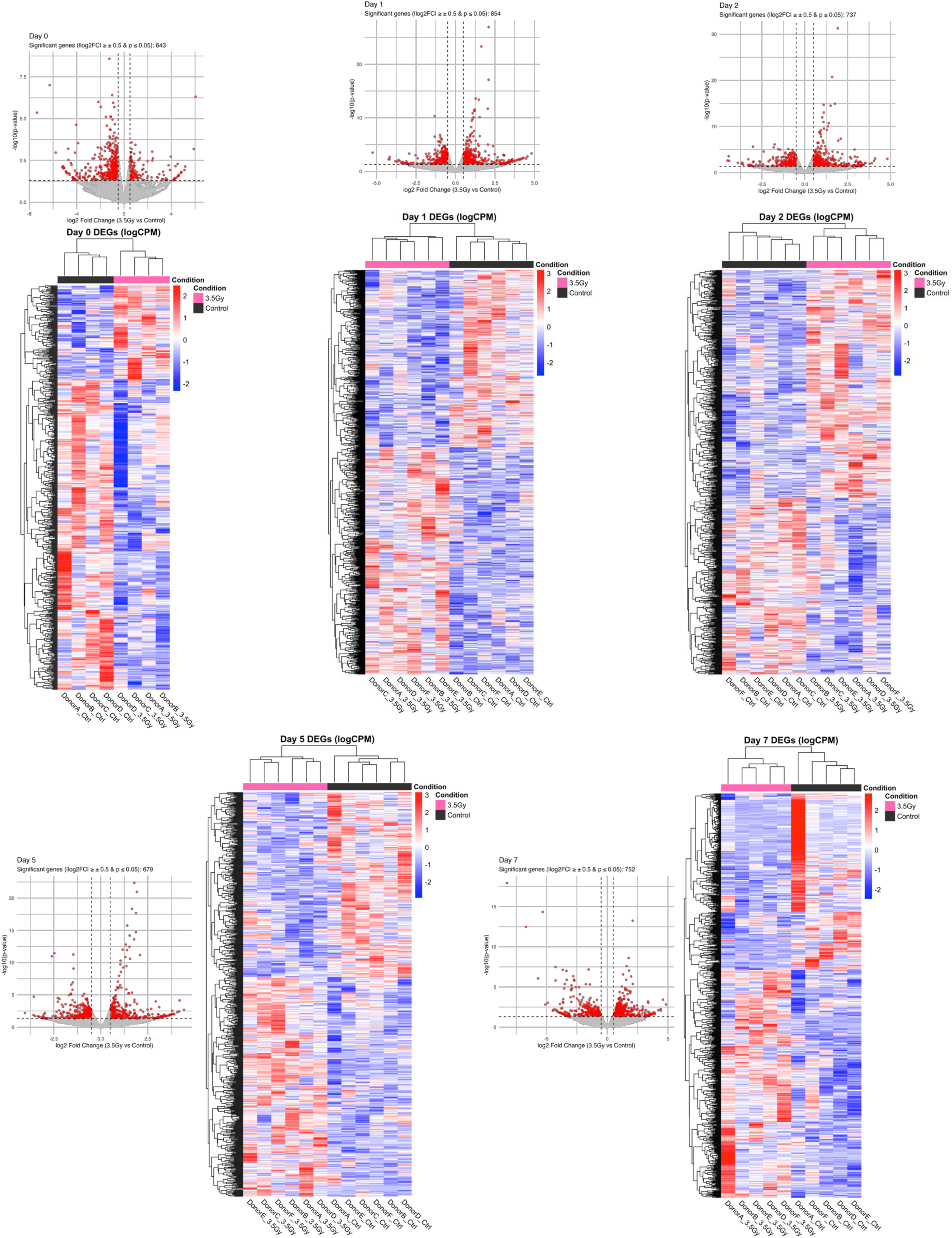
Differentially expressed genes in irradiated vs. control skin by donor across time points. Volcano plots display the results of differential gene expression analysis for each time point following 3.5 Gy-radiation exposure compared to control. Each point represents a gene, with the x-axis showing the log2FC and y-axis showing the –log10 p-value. Genes highlighted in red are those that met the significance thresholds of an absolute log_2_FC ≥ ± 0.5 and p-value ≤ 0.05. The number of significant genes (meeting both thresholds) is indicated within each volcano plot. Heatmaps are visualizing the expression patterns of these significant genes across individual donors at each corresponding time point. Each row in the heatmap represents a gene, and each column represents an individual sample from a unique donor at that time point. The color intensity corresponds to the gene’s expression level (red = increased, blue = decreased).

**Fig. S2.**
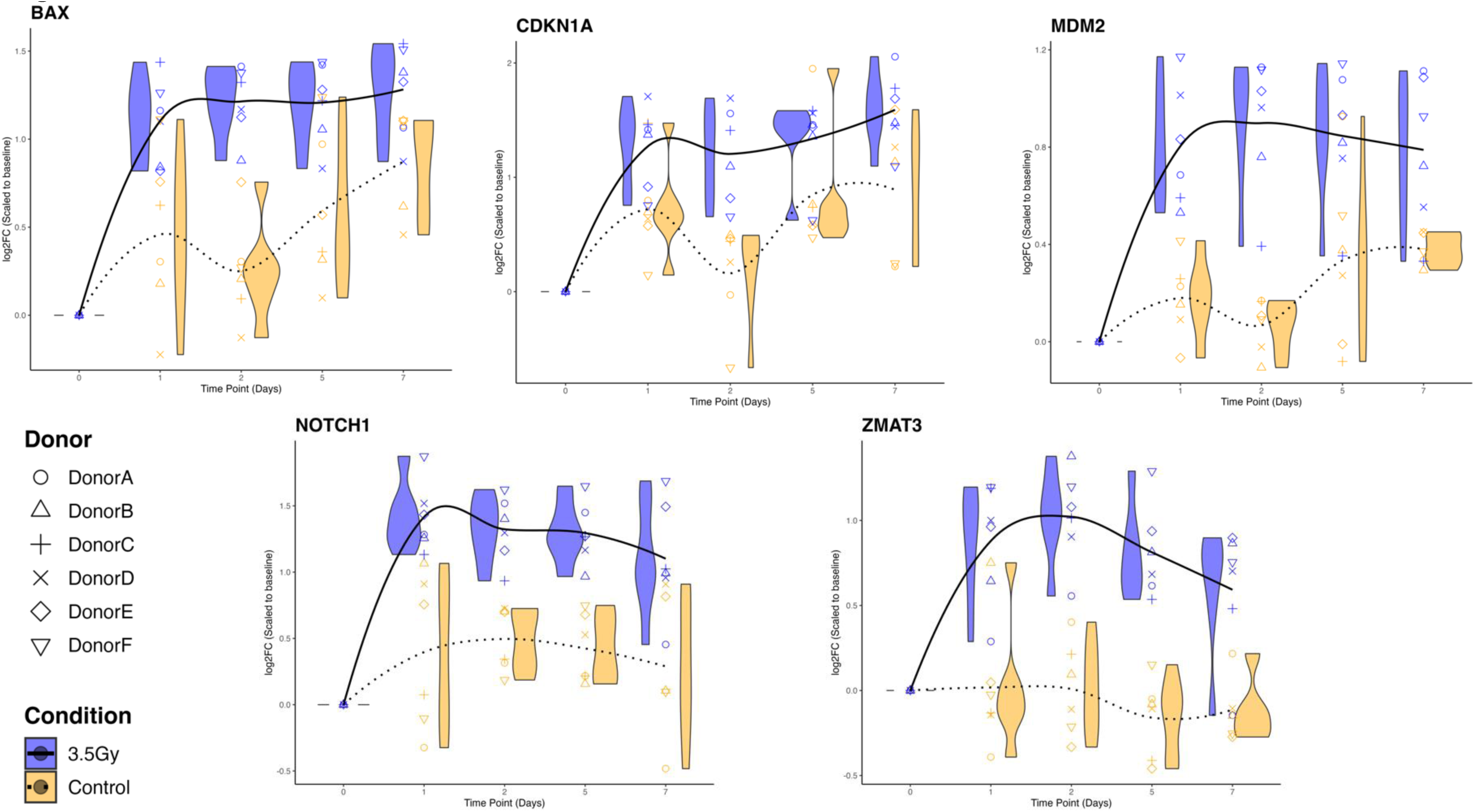
Trajectories of p53 target genes over time following irradiation. Trajectory plots illustrate the temporal expression patterns of specific genes following exposure to 3.5 Gy-irradiation (blue) or control (yellow). Each point represents the log_2_ fold change (log_2_FC) in gene expression after being scaled to the earliest time point (day 0). Individual donors are distinguished by unique symbols. The x-axis represents time, and the y-axis displays the log_2_FC. Violin plots are overlaid at each time point to show the distribution of log_2_FC values across all donors within each treatment group. Solid and dotted lines represent the mean log_2_FC across all donors within the 3.5 Gy and control groups, respectively, at each time point.

**Table S1.** Pro-fibrotic pathways and upstream regulators. Table containing the number of downstream target genes, z-score and Benjamini-Hochberg (BH)-corrected p-value for days 2 and 7, corresponding to **Figure 4A**.

| Pathway / Process / Regulator | Number of downstream gene targets | Z-score | BH-corrected p-value | Number of downstream gene targets | Z-score | BH-corrected p-value |
| --- | --- | --- | --- | --- | --- | --- |
|  | Day 2 |  |  | Day 7 |  |  |
| Pulmonary Fibrosis Idiopathic Signaling Pathway | 17 | 0.775 | 2.32319035 | 24 | 1.877 | 1.41237172 |
| Extracellular matrix organization | 11 | 0.302 | 4.96460698 | 9 | 1 | 1.16210297 |
| Assembly of collagen fibrils | 6 | 1.633 | 1.1197002 | 5 | 1.342 | 3.0962585 |
| Hepatic fibrosis | 21 | 0 | 1.20125381 | 12 | 0 | 0.83619627 |
| SMAD3 | 25 | 2.095 | 3.61730163 | 26 | 2.568 | 3.82421016 |
| SMAD4 | 35 | 2.582 | 5.78144247 | 45 | 2.919 | 10.7612484 |
| SNAI1 | 15 | 1.143 | 2.70839086 | 16 | 0.387 | 2.94322408 |
| SNAI2 | 0 | 0 | 0 | 11 | 1.953 | 2.05403718 |
| TGF beta | 36 | 0.438 | 5.33313354 | 40 | 1.976 | 6.92916043 |
| TGFB1 | 113 | 2.078 | 8.69939088 | 144 | 3.479 | 19.4325154 |
| TGFB2 | 18 | 1.468 | 3.4465511 | 24 | 1.334 | 6.47395085 |
| TGFBR1 | 10 | 1.177 | 1.9251388 | 11 | 1.16 | 2.14855399 |
| TGFBR2 | 32 | 1.2 | 4.85062614 | 32 | 2.664 | 4.69676627 |
| TWIST1 | 18 | 0.983 | 3.20171389 | 23 | 0.901 | 5.41851824 |
| Fibrogenesis | 43 | -0.543 | 3.47269136 | 60 | 2.162 | 9.66812034 |
| Fibrosis | 98 | 0.143 | 8.22160462 | 104 | 0.596 | 9.58781214 |

**Table S2.**
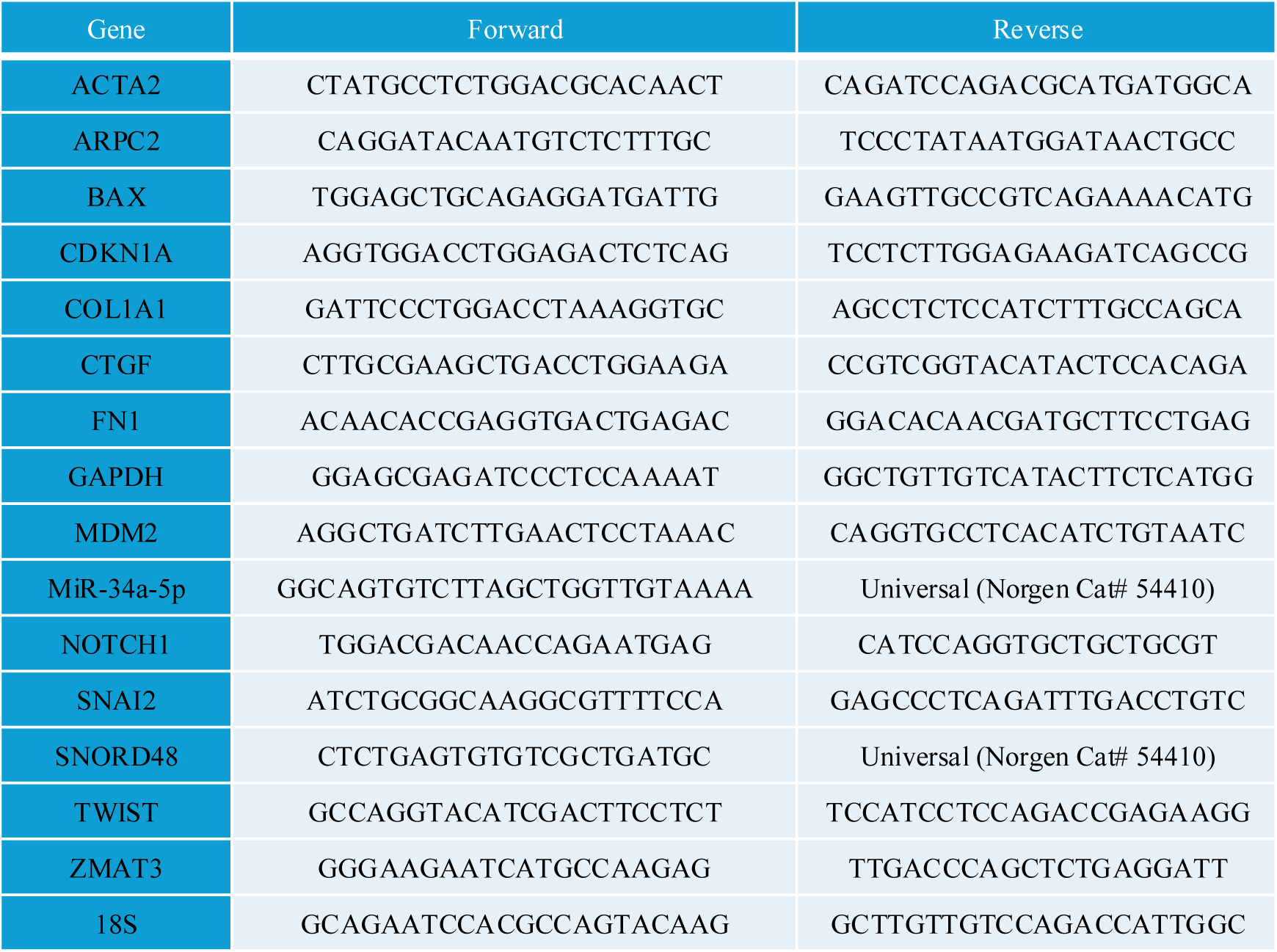
Human primer sequences used for qPCR.

**Table S3.** Primary and secondary antibodies utilized for immunohistochemistry, Western blotting and immunofluorescence.

| Antibody | Manufacturer | Catalogue # | Method Used | Dilution |
| --- | --- | --- | --- | --- |
| **E-Cadherin** | Cell Signaling Technology (Danvers, MA, USA) | 3195 | IHC | 1:200 |
| **E-Cadherin** | Cell Signaling Technology (Danvers, MA, USA) | 3195 | WB | 1:1000 |
| **Vimentin** | Cell Signaling Technology (Danvers, MA, USA) | 5741 | IHC | 1:200 |
| **γH2AX** | Cell Signaling Technology (Danvers, MA, USA) | 9718 | IF | 1:500 |
| **MDM2** | Invitrogen (Carlsbad, CA, USA) | PA5-13008 | IF | 1:25 |
| **GAPDH** | Cell Signaling Technology (Danvers, MA, USA) | 2118 | WB | 1:1000 |
| **Goat anti-Rabbit IgG (H+L) Alexa Fluor 488** | Invitrogen (Carlsbad, CA, USA) | A-11008 | IF Secondary | 1:500 |
| **Goat anti-Mouse IgG (H+L) Alexa Fluor 594** | Invitrogen (Carlsbad, CA, USA) | A-11005 | IF Secondary | 1:500 |

## Notes

### Competing Interest Statement

The authors have declared no competing interest.

